# Whole-Slide Mapping of Tumor Tissue Fiber Architecture via Computational Scattered Light Imaging

**DOI:** 10.64898/2026.02.23.707464

**Authors:** Hamed Abbasi, Loes Ettema, Rens Van Elk, Meike Eskes, Michail Doukas, Sjors A. Koppes, Stijn Keereweer, Miriam Menzel

## Abstract

Mapping peritumoral collagen fiber directionality in solid tumors may assist in determining cancer progression and support more personalized prognoses. However, existing microscopy techniques are often limited by a restricted field of view, high cost, or incompatibility with paraffin-treated tissues. Computational scattered light imaging (ComSLI) is a cost-effective whole-slide microscopy technique that reveals fiber orientations independent of sample preparation. Using glioma, colorectal, and head and neck cancer samples, we show for the first time that ComSLI maps fiber orientations in paraffin-treated tumor tissues, visualizes tumor growth pathways and desmoplastic reactions, and allows the study of collagen orientations relative to tumor boundaries.

## 1. Introduction

Cancer is a leading cause of death worldwide, with solid tumors accounting for the majority of cases [1]. For most patients diagnosed with a solid tumor, surgery remains the primary treatment option, often followed by postoperative adjuvant therapy. The choice of the appropriate postoperative treatment depends on the risk of disease recurrence and metastasis. Risk stratification in oncology enables a more personalized approach to patient management by integrating biological, anatomical, and molecular information to estimate the probability of disease recurrence or progression. By distinguishing between low-, intermediate-, and high-risk patients, clinicians can tailor treatment intensity, avoid overtreatment in those with indolent disease, and focus aggressive interventions on individuals most likely to benefit. This not only optimizes therapeutic efficacy and resource allocation but also improves patient quality of life and long-term outcomes. The risk of disease recurrence and metastasis is determined by pathologists through the assessment of various biomarkers on microscopy slides of the resected tumor and lymph nodes. For example, in oral squamous cell carcinoma (OSCC), the risk of recurrence and metastasis is categorized according to the worst pattern of invasion (WPOI): scales 1–3 indicate low invasiveness, while scales 4–5 indicate high invasiveness. Although WPOI scoring plays a key role in treatment guidelines, it is manually determined through visual inspection of histology images and is subject to significant inter- and intra-observer variability [2-6]. Besides WPOI, depth of invasion (DOI) is also an essential factor in determining whether a patient might need additional treatment, such as an elective neck dissection (END), as it can be an indicator of occult lymph node metastasis. However, about 80% of the patients who receive END have no occult lymph node metastasis, and END can be associated with several adverse effects, such as edema, pain, neck and shoulder discomfort, as well as shoulder disability [7]. Hence, there is a need for an objective method to determine the risk of recurrence and metastasis more accurately to provide more tailored treatments for patients with cancer and to avoid both under- and over-treatment.

Recent studies have identified a promising prognostic risk factor for regional and distant metastasis: changes in the organization and directionality of (collagen) fibers in the extracellular matrix (ECM) [8, 9]. There is a dynamic interplay between peritumoral fibers and tumor evolution [10]. In an aggressive tumor, metastatic tumor cells interact with oriented ECM fibers and invade the basement membrane and vessels [11, 12]. Remodeling of the ECM aligns the collagen fibers from parallel to perpendicular to the tumor boundary, thereby opening passages for tumor cell migration [13, 14]. This transition is described as the shift from tumor-associated collagen signature 2 (TACS-2) to TACS-3 [15], although other studies have found a higher number of different TACS patterns [16]. Figure 1 illustrates the evolution of collagen fiber organization during tumor progression.

**Fig. 1.**
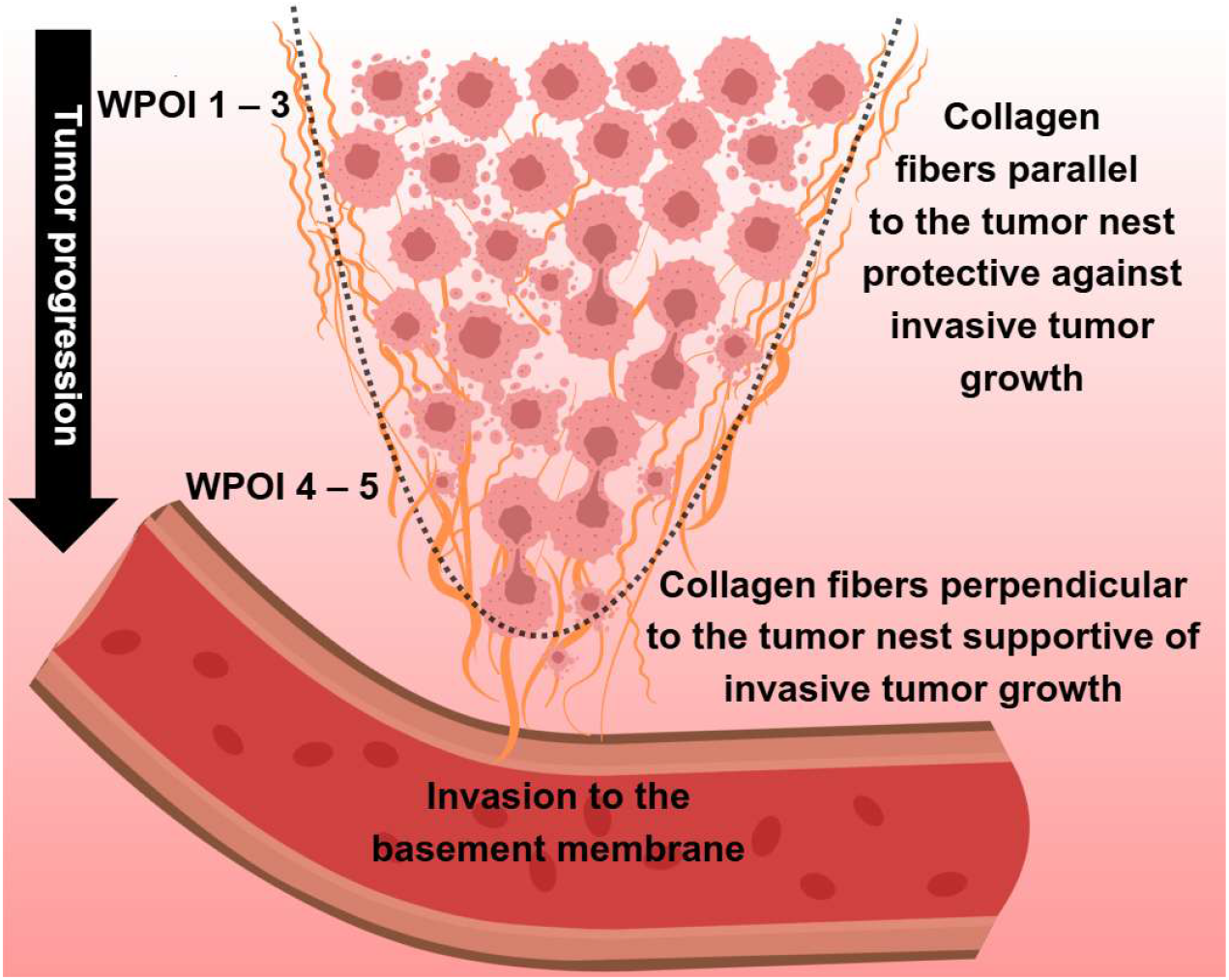
Evolution of collagen fiber organization during tumor progression. In low-score WPOI (1-3), a dense collagen matrix surrounds the tumor cells, whereas in high-score WPOI (4-5), metastatic cells interact with perpendicularly oriented collagen fibers and invade the tumor basement membrane.

Although the effects of the mechanical properties of the microenvironment on tumor cell phenotypes are still being elucidated [17, 18], several retrospective studies have observed correlations between prognosis and collagen fiber properties, including elongation, thickness, straightness, density, and orientation [14, 19]. Such differences in collagen organization between aggressive and non-aggressive tumors have been reported in the breast [16, 20-27], gastric [28], hepatic [29], lung [30], ovarian [31, 32], head & neck [33-35], esophageal [9], renal [36], pancreatic [37-43], glioblastoma [44], bladder [45], prostate [46-48] and colorectal cancers [24, 49]. These studies showed that oriented collagen fibers guide tumor cells into surrounding tissues. In non-aggressive tumors, a dense collagen matrix surrounds the tumor nest, whereas in aggressive tumors, collagen fibers perpendicular to the tumor boundary predominate. This reorganization facilitates directional migration of cancer cells into the surrounding stroma, guiding them toward blood and lymphatic vessels and enabling intravasation [22]. Perpendicular fiber alignment in peritumoral regions has been consistently associated with more aggressive disease, higher recurrence rates, and reduced overall survival across multiple tumor types [50, 51].

Over the past two decades, extensive research has focused on collagen reorganization parameters, including fiber angle and alignment. These parameters are increasingly recognized as stromal biomarkers linked to disease progression and survival in solid tumors. However, the supporting evidence is stronger in some cancers, such as breast and pancreatic cancer, than in others, like head and neck cancer, which remain less explored. While standard bright-field microscopy of hematoxylin and eosin (H&E)-stained sections provides crucial histological information and is commonly used by pathologists to determine the tumor boundary, it does not allow objective or precise quantification of collagen fiber directionality. Therefore, different imaging techniques, mainly optical, were developed to visualize fiber orientation in tissues [52, 53]. Over the years, these methods have demonstrated that collagen fiber orientation relative to malignant epithelium carries prognostic information across a variety of tissues. Nonetheless, little effort has been made to make such biomarkers accessible to clinicians without requiring expensive imaging systems or significant workflow changes [42, 54]. While most of the literature focuses on changes in collagen fiber orientation, studying orientation changes in other fiber types, such as muscle or nerve fibers, can also be valuable for understanding how tumor growth affects tissue architecture, for example when visualizing tumor growth pathways in muscle tissue or reorganization of nerve fibers in brain cancer samples.

There exist techniques that allow to indirectly estimate the orientation of different fiber types through image processing, such as bright-field microscopy [26, 27], optical coherence tomography (OCT) [55], and second harmonic generation (SHG) [24, 29, 31, 32, 47]. Other techniques can directly measure the local fiber orientation, like diffusion magnetic resonance imaging (dMRI) [56], small-angle X-ray scattering (SAXS) [56], polarimetric SHG (pSHG) [57], polarization-sensitive OCT (PS-OCT) [58], polarized light imaging (PLI) [59], and polychromatic polarization microscopy (PPM) [42, 43]. However, each of these methods has limitations: dMRI has poor resolution and requires an expensive MRI scanner; SAXS and (p)SHG have a limited field of view (FoV); SAXS is time-consuming and requires costly synchrotron light sources; (p)SHG requires non-centrosymmetric molecular structures and is not suited for visualizing nerve fibers; PS-OCT, PLI, and PPM cannot correctly determine the individual orientations of multiple crossing or out-of-plane fibers, and they rely on birefringence-preserving sample preparations [60] while PPM also requires individual calibration for different stainings to compensate for the wavelength dependency of the birefringence across different stains [43]. The requirement of sufficient birefringence contrast makes polarization-based techniques incompatible with common sample preparations. For example, permeabilization with Triton X-100, a reagent commonly used in immunohistochemistry fluorescence staining, impairs the birefringence of myelinated nerve fibers by solubilizing the lipid bilayers of the surrounding myelin sheath [61]. Previously, we also showed that formalin-fixed paraffin-embedded (FFPE) brain samples, which were treated with alcohol and xylene before paraffin embedding, have insufficient birefringence contrast to determine nerve fiber orientations in the white matter with polarization microscopy [62]. Compatibility with FFPE tissue is particularly important as FFPE is the standard preservation method in clinical pathology and enables access to large archival cohorts.

Computational scattered light imaging (ComSLI) fills this gap by offering a label-free, staining-independent technique applicable to various sample preparations, including FFPE tissues [57, 62, 63]. Instead of relying on birefringence, ComSLI uses anisotropic light scattering to disentangle fiber architecture without the need for time-consuming raster scanning, enabling high-resolution, large-scale imaging of interwoven biological fiber networks. Requiring only an LED light and a camera, it provides a fast, cost-effective, and objective method for quantitatively measuring the orientation of fibers in microscopy slides. While techniques like SHG are restricted to specific fiber types, ComSLI can determine the orientation of all kinds of directed structures, including collagen, muscle, nerve, and elastin fibers [62]; to identify the specific fiber type being imaged, staining or anatomical knowledge is needed. With a much higher resolution than dMRI, a larger field-of-view than SAXS or pSHG, more fiber orientations per pixel detected than polarization-based techniques, a simpler and more cost-effective setup than all other techniques, and the ability to visualize all kinds of fibers in variously prepared tissue samples, ComSLI is a highly promising technique for analyzing microscopic fiber organization in cancer tissues.

However, up to now, ComSLI has only been applied to investigate fiber architecture in non-tumor tissues, particularly nerve fibers in the brain [56, 64, 65]. In this study, we extend its application to visualize fiber structures across a range of solid tumor types, showing for the first time the use of ComSLI in diseased non-brain tissue samples. We apply ComSLI to FFPE sections of glioma, colorectal, and head and neck cancer, compare its performance to polarization-based microscopy, study its ability to visualize tumor growth pathways and desmoplastic reactions, and analyze the orientation of collagen fibers relative to the tumor invasive front in samples with different WPOIs.

## 2. Material and Methods

### 2.1 Ethics statement

#### Human tissue

The human tissue sections were obtained from a tissue archive at Erasmus Medical Center (Erasmus MC Cancer Institute), Rotterdam, the Netherlands, approved by the Medisch Ethische Toetsings Commissie (METC) under number MEC-2023-0587. The study was conducted in full compliance with the principles of the Declaration of Helsinki of 1975, the International Council for Harmonisation’s Good Clinical Practice (ICH GCP) guidelines, and the laws and regulations of the Netherlands. The study was also evaluated and approved by the Human Research Ethics Committee (HREC) of Delft University of Technology (TU Delft), Delft, the Netherlands, and a Data and Material Transfer Agreement was obtained between Erasmus MC and TU Delft. The tissue samples were obtained from patients during oncological surgery as standard clinical procedure, and no extra tissue was removed for this study. The tissue slides were returned to the archive after measurements.

#### Animal tissue

The rat brain used in this study was obtained from a healthy male Wistar rat (3 months old). All animal procedures were approved by the institutional animal welfare committee at Forschungszentrum Jülich GmbH, Germany, and were in accordance with the European Union and National Institutes of Health guidelines for the use and care of laboratory animals, as well as the ARRIVE guidelines. Euthanasia of the rat was carried out under controlled isoflurane inhalation followed by decapitation. The mouse brain samples were borrowed from different studies [66, 67] which were conducted according to the guidelines of the Declaration of Helsinki and approved by the Institutional Review Board (and Animal Ethics Committee) of Erasmus MC, Rotterdam, the Netherlands, under the approved work protocols SP2300047, SP2300149 and SP2300238, approval date 08–05–2023, 12–06–2024 and 29–02–2024 respectively, covered by the national project license CCD number AVD101002017867. The mice were housed and treated under the animal protocols approved by the Ethical Committee of the Animal Welfare Body of Erasmus Medical Center, the Netherlands. Experiments were carried out according to ARRIVE guidelines [66, 67].

### 2.2 Sample preparation

#### Human tissue

The human tissue samples were obtained from patients during oncological surgery and prepared directly afterwards. The mandible sample containing bone was first decalcified using DecalMATE by Milestone Medical. Afterwards, all samples were fixed in 4% formaldehyde for 24–72 hours, dehydrated in increasing alcohol series (70%, 80%, 90%, 96%, 100% ethanol), treated with xylene, embedded in paraffin, and cut with a microtome (Leica RM2165) into 4-μm-thin sections. The sections were placed in a decreasing alcohol series to remove the paraffin, mounted on glass slides, stained with hematoxylin and eosin (H&E), or picrosirius red (PSR), and then cover-slipped with a mounting medium (HE 600 Coverslip Activator, Roche).

#### Animal tissue

Rat brain sample: The brain was removed from the skull within 24 hours after death, fixed with 4% buffered formaldehyde for several weeks, cryo-protected with 20% glycerin and 2% dimethyl sulfoxide, deeply frozen, and coronally cut with a cryostat microtome (Polycut CM 3500, Leica, Microsystems, Germany) into sections of 60 μm. Section no. 28 was selected for evaluation. The section was mounted on a glass slide, embedded in 20% glycerin solution, cover-slipped, and sealed with lacquer.

Mouse brain samples: The samples with glioma were borrowed from different studies. a NOD/SCID mouse (male, 10-12 weeks old) with orthotopic patient-derived glioblastoma xenograft tumor in the right hemisphere, where the mouse was sacrificed at 28 days post-inoculation of GS756 cell line [66] (shown in Fig. 2f–m) and an NMRI nude mouse (male, 6– 7 weeks old) bearing a tumor in the right hemisphere, where the mouse was sacrificed at 14 days post-inoculation of U87-MG cell line [67] (shown in Fig. S2). The sections are FFPE H&E-stained 4 μm sections prepared as described for the human tissue.

**Fig. 2.**
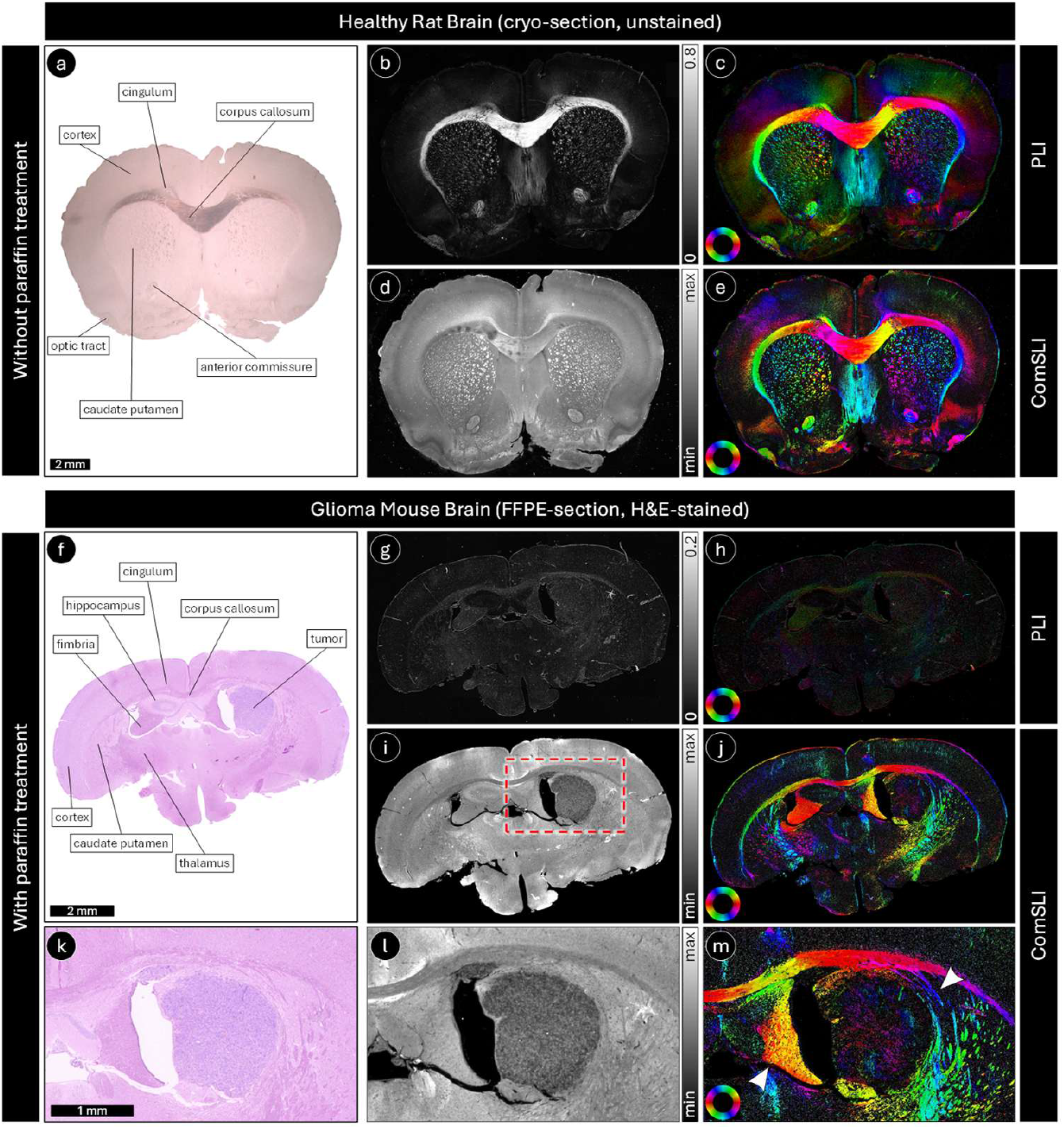
Polarized light imaging (PLI) versus ComSLI in healthy and glioma brain samples without and with paraffin treatment. Top: The figure shows PLI and ComSLI measurements in an unstained, 60-μm-thin cryo-section of a healthy rat brain (without paraffin treatment). Bottom: The figure shows PLI and ComSLI measurements in an H&E-stained, 4-μm-thin FFPE section of a mouse brain with glioma (with paraffin treatment). (a) Bright-field digital microscopy image of the healthy rat brain section, captured with slightly oblique illumination to enhance the contrast. Anatomical regions are labeled exemplarily for the left hemisphere. (f) Bright-field digital microscopy image of the mouse brain section with glioma. Anatomical regions are labeled exemplarily in the left hemisphere, as well as the tumor in the right hemisphere (hippocampus). (b) and (g) PLI retardance maps. (c) and (h) PLI fiber orientation maps. (d) and (i) ComSLI average scattering maps. (e) and (j) ComSLI fiber orientation maps. (k), (l), and (m) show the zoom-in of (f), (i), and (j), respectively (region marked by the red dashed rectangle in (i)). The arrowheads in panel (m) highlight regions with altered fiber orientation resulting from tumor growth in the right hemisphere. The upper arrowhead indicates nerve fibers surrounding the tumor, while the lower arrowhead points to the deformed fimbria. Fiber orientation maps of both PLI and ComSLI are color-coded according to the respective color wheel in the bottom left. While PLI can determine nerve fiber orientations in sections not treated with paraffin, similar to ComSLI, it cannot retrieve the orientations in FFPE sections because the birefringence is impaired during the preparation process, which leads to low retardance. In (g) and (h), the brightness was intentionally enhanced so that the structures become better visible. ComSLI, in contrast, can retrieve fiber orientations regardless of the sample preparation.

### 2.3 Bright-field microscopy and histopathological annotations

The stained microscopy slides were scanned using the Nanozoomer 2.0 HT digital slide scanner (Hamamatsu Photonics K.K.), offering a 20X magnification and a pixel size of 0.46 μm. Bright-field microscopy images of H&E-stained sections were used to annotate the tumor and fibrosis regions in the microscopy slides by experienced pathologists blinded to the ComSLI or PLI microscopy images. The unstained rat brain microscopy slide was scanned using the Keyence VHX-6000 Digital Microscope (with VH-ZST objective, 2X). For the unstained sample, slightly oblique illumination was used to enhance the contrast.

### 2.4 PLI

PLI measurements were performed using a customized polarization microscope (CM502-532, Cerna Birefringence Imaging Microscope, by Thorlabs), which can measure sample retardance and azimuths. The azimuth, corresponding to the fast axis orientation, is determined from 0° to 180°. The sample is illuminated by green, linearly polarized light with different orientations {0°, 45°, 90°, 135°}, and the transmitted light passes through a circular analyzer before being recorded by a camera. The microscope is equipped with a green LED light source (M530L4, Thorlabs), a 532 nm bandpass filter with a full width at half maximum (FWHM) of 1 nm, and liquid crystal modules. The camera is a monochrome CMOS with 2448 × 2048 pixels and a 3.45 μm pitch (imaging area of 8.4456 mm × 7.0656 mm). A 4X microscope objective (N4X-PF, Thorlabs) with an effective focal length (EFL) of 50 mm, working distance of 17.2 mm, and a numerical aperture (NA) of 0.13 was used in this study, resulting in a pixel size of 0.86 μm in object space and a field of view (FoV) of 2.1 mm × 1.8 mm. The dataset measured here consists of four images acquired with linearly polarized ingoing light with 0°, 45°, 90°, and 135° direction of polarization. An in-house developed Python script was used to calculate for each image pixel the Fourier coefficients of the resulting sinusoidal signal, and the retardance and fast axis orientation (in-plane fiber orientation) were computed from the amplitude and the phase of the signal, respectively, as described in [68, 69].

### 2.5 Experimental setup of ComSLI

The imaging part of the ComSLI experimental setup consists of a monochrome CMOS camera (BASLER acA5472-17um, 20 Megapixels) with a 5472 × 3648 pixel resolution and a 2.4 μm pixel pitch, attached to a camera lens with a focal length of 120 mm and a focal ratio of 5.6 (Rodenstock Apo-Rodagon-D120) using a 148 mm long extension tube. This camera and lens combination provides a FoV of 1.6 cm × 1.1 cm, and an optical resolution between 3.91 μm and 4.38 μm as measured by a USAF target.

A fiber-coupled white-light LED (Prizmatix) was used to illuminate the samples placed at a 45-degree polar angle. The light source emits with a peak wavelength of 443 nm (range 400– 750 nm); light was guided through a 2-meter-long multimode fiber with a 1500 μm core diameter, and passed through a collimator (FCM1-0.5-CN, Prizmatix) and an engineered diffuser (ED1-S20-MD, Thorlabs) for beam shaping before reaching the sample. For each sample, 24 images (each being the average of 4 images) were acquired by performing angular (azimuthal) measurements and measuring intensities discretely over a full circle (every 15 degrees). The exposure time was individually adjusted for each sample, typically in the range of tens of milliseconds, to balance a broad dynamic range while avoiding saturation. Figure S1 presents a schematic of the ComSLI setup in which a collimated light beam obliquely illuminates a fiber bundle in the sample and scatters mainly in directions perpendicular to the fiber axis. This scattering pattern produces peak pairs in the azimuthal intensity profile, *I*(*φ*), with the midpoint between the peaks corresponding to the in-plane fiber orientation.

### 2.6 ComSLI data processing

The intensity images acquired with ComSLI were processed with the scattered light imaging toolbox (SLIX) [70] to obtain different parameter maps. For each image pixel, SLIX constructs an azimuthal intensity profile from images acquired at different illumination angles and determines the in-plane fiber orientations by detecting prominent peaks in this profile, corresponding to dominant scattering directions: For each peak pair (two prominent peaks lying 180°±35° apart), the in-plane fiber orientation is determined by the mid-position between the two peaks. If a line profile contains a single prominent peak, the fiber orientation is given by the position of the peak. Profiles with 3, 5, or more than 6 peaks are not evaluated. SLIX provides an average intensity map, which shows the average signal intensity across different illumination angles, and a fiber orientation map (FOM), which shows the in-plane fiber orientations in different colors according to a rainbow-color wheel. Fiber orientations are indicated as azimuth angles and range from 0° to 180° (moving anti-clockwise toward larger values). The angles shown in the original FOMs created by SLIX are represented as absolute angles. To study and analyze fiber orientations relative to the tumor boundary, an in-house Python script was developed to express the fiber orientations at each pixel with respect to the tumor invasive front, thus producing relative FOMs. An in-house MATLAB script was used to generate relative FOMs in which fibers that are (semi-)parallel or (semi-)perpendicular to the tumor invasive front are shown in green or magenta, respectively. The FOMs were masked with the average intensity map to remove the background signal that had a low average scattering intensity.

## 3. Results

Figure 2 illustrates how ComSLI performs in comparison to PLI when being applied to healthy and glioma brain tissue prepared without and with paraffin treatment (here shown for a coronal rat and mouse brain section, respectively). In cryo-brain sections not treated with paraffin, both techniques reveal the expected tissue anisotropy, and the fiber orientation patterns derived from ComSLI align well with those obtained from PLI (cf. Fig. 2c and e). However, once the brain tissue has undergone paraffin embedding and staining, the birefringence signal required for PLI is markedly diminished (compare the retardance in corpus callosum in Fig. 2b and g). This results in poor contrast in the retardance and orientation maps (see Fig. 2g and h), making the method unsuitable for visualizing nerve fiber organization in such samples. In contrast, ComSLI continues to detect scattering anisotropy and provides consistent fiber orientation information also in FFPE brain sections (see Fig. 2j). These results demonstrate that ComSLI is robust to fixation, paraffin, and alcohol treatment, as well as staining, which enables orientation mapping in a broader range of histological preparations than is possible with PLI. ComSLI is also able to visualize the orientations of the nerve fibers surrounding the tumor (annotated in Fig. 2f), as well as altered fiber orientations resulting from tumor growth in the right hemisphere (hippocampus), highlighted by arrowheads in Fig. 2m. The hippocampus is not visible anymore in the right hemisphere, the fimbria is significantly deformed (bottom arrowhead), and fibers from the caudate putamen are surrounding the tumor (top arrowhead). An additional glioma sample is shown in Supplementary Fig. S2. In both glioma samples, the tumor region appears significantly darker in the average scattering maps than the surrounding tissue areas, enabling a clear delineation of the tumor borders (Fig. 2i, l and S2b). Within the tumor areas, only a few directed structures are visible (Fig. 2m and S2c), indicating that no fibrillar collagen is present within the glioma. The observed fiber patterns reveal changes in local fiber alignment at the tumor boundary, which may reflect tumor-induced tissue remodeling, displacement, or infiltration of adjacent neural structures. Such information on peritumoral fiber organization is difficult to obtain with conventional histology alone and highlights the potential of ComSLI to provide complementary structural insight into tumor– tissue interactions.

Figure 3 shows the potential of ComSLI to characterize structural changes associated with tumor invasion. The human rectal cancer specimen includes a region where the tumor (annotated with a green line in the H&E digital scan in Fig. 3a) advances into the surrounding stromal tissue (zoom-in shown in Fig. 3d). In the ComSLI average scattering and orientation maps (Fig. 3e and f), the area along the invasive margin exhibits a marked reorganization of the fibrous network (arrowhead in Fig. 3f). The trajectories of the newly aligned fibers correspond to the direction in which the tumor expands (dotted purple arrow in Fig. 3e), which supports the idea that stromal remodeling accompanies invasion. The method also highlights the presence of fibrosis (annotated with yellow lines in Fig. 3a), where the fiber arrangement differs noticeably from that of the adjacent tissue (arrowhead in Fig. 3c). The distinction between preexisting and newly deposited fibers becomes apparent through their differing orientations, suggesting that ComSLI can capture the dynamic remodeling processes that accompany tumor progression.

**Fig. 3.**
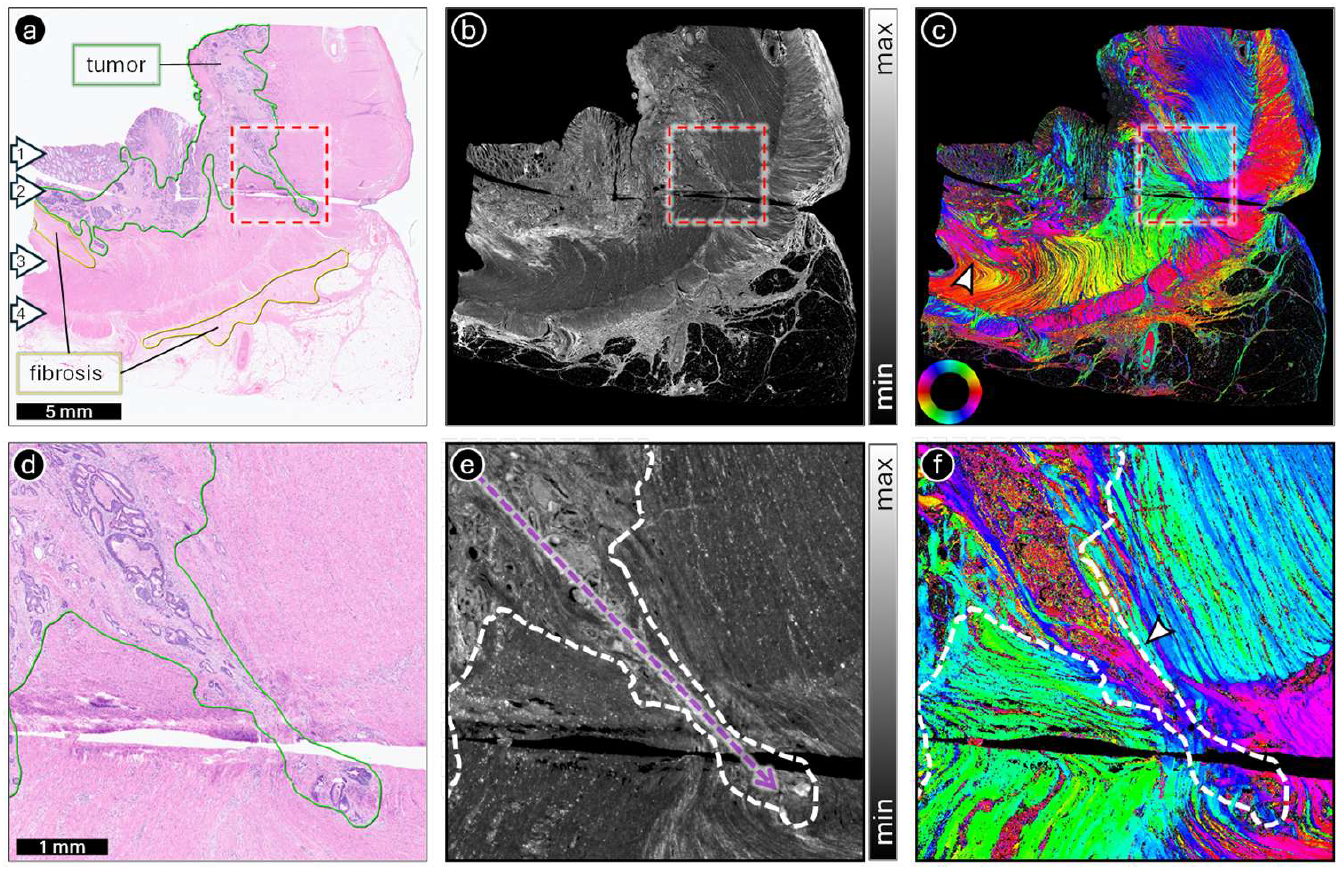
ComSLI for visualizing the tumor growth path. The figure shows the ComSLI measurement of a human rectal cancer specimen (H&E-stained, 4-μm-thin FFPE section). (a) Bright-field digital scan. The arrows indicate (1) mucosa with epithelium, (2) submucosa, (3) circular muscle layer, and (4) longitudinal muscle layer. Tumor and fibrosis regions are annotated in green and yellow, respectively. The dashed red square indicates an area at the invasive front where the tumor has grown further toward the rectal wall. (b) Average scattering map. (c) Fiber orientation map, color-coded according to the shown color wheel. The white arrowhead points towards one of the fibrosis areas. (d-f) Zoom-ins on the dashed red squares in (a-c). The dashed arrow in (e) and the arrowhead in (f) indicate the tumor growth pathway. The fiber orientation map clearly highlights the different fiber orientations along the tumor growth pathway. The area with fibrosis also shows a distinct fiber orientation compared to neighboring fibers (see arrowhead in c), highlighting the newly generated fibers that are replacing the preexisting ones.

Figure 4 further explores the utility of ComSLI in visualizing tumor-associated tissue responses, here in a case of human oral cavity cancer that extends from soft tissue into the mandible. As the tumor penetrates the cortical layer and spreads into trabecular bone, the stromal reaction becomes evident. The tumor originated in soft tissue surrounding the cortical bone of the mandible and, as it progressed, invaded the cortical bone before extending into the trabecular bone. The original location of the cortical bone is indicated by a white dotted line in Fig. 4. The fiber orientation patterns (Fig. 4c) reveal fiber orientations in the tumor region (annotated by a solid black line in Fig. 4a) that deviate from those in unaffected regions (the deviating fiber pathways are indicated by white arrowheads in Fig. 4c). This shift reflects the desmoplastic reaction (present all over the tumor region), in which fibroblasts reorganize the extracellular matrix and generate new collagen fibers as part of local tissue defense. The changes in fiber orientation in these areas allow ComSLI to delineate regions of active remodeling and bone degradation, demonstrating its effectiveness in capturing the complexity of tumor–stroma interactions.

**Fig. 4.**
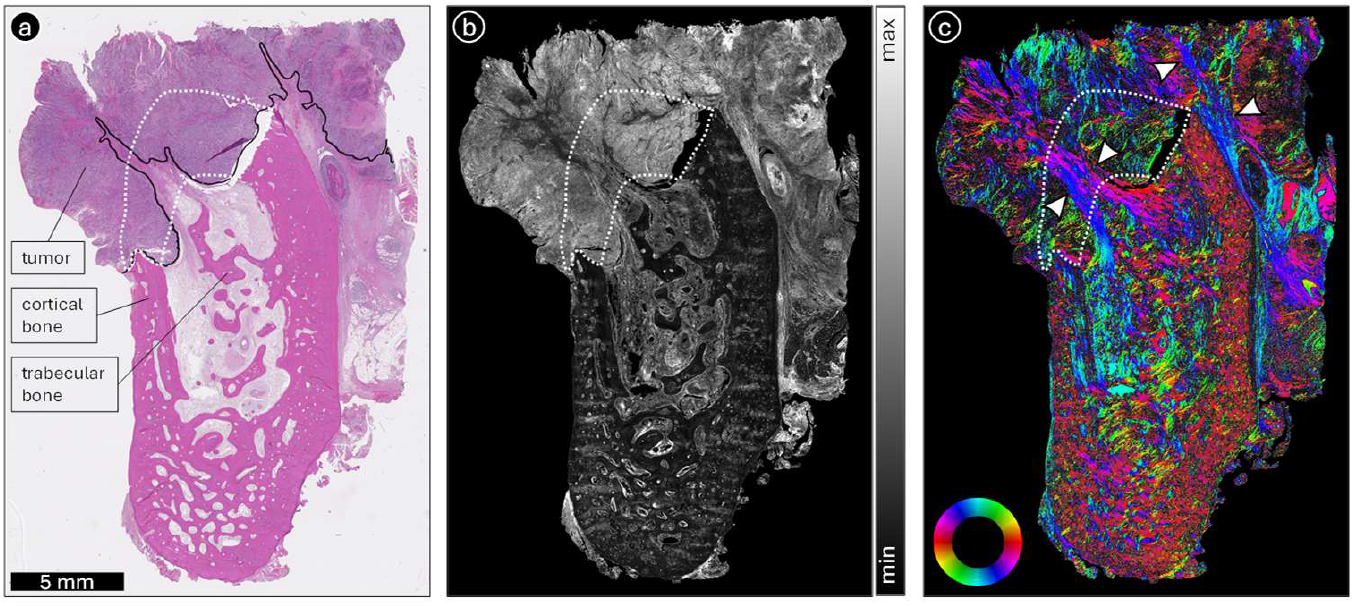
ComSLI for visualizing desmoplastic reactions. The figure shows the ComSLI measurement of an human oral cavity cancer specimen (H&E-stained, 4-μm-thin FFPE section). (a) Bright-field digital scan, with cortical and trabecular bone labeled, and the tumor invasive front annotated by a solid black line. The area where the cortical bone was originally located is annotated by a white dotted line. (b) Average scattering map. (c) Fiber orientation map, color-coded according to the shown color wheel. The tumor developed in soft tissue surrounding the cortical bone of the mandible, and as it expanded, it invaded the cortical bone and then extended into the trabecular bone. In response to the cancer, fibroblasts contribute to bone degradation and produce new collagen fibers that differ in orientation from the preexisting fibers, a process known as the desmoplastic reaction, which is present throughout the tumor region in this section. The fiber orientation map in (c) clearly highlights the distinct fiber orientations in regions with desmoplastic reaction, indicated by white arrowheads.

Figure 5 illustrates how ComSLI can be used to assess collagen fiber organization in relation to the tumor invasive front in human oral cavity cancer. Representative examples are shown for tumors with distinct patterns of invasion, including a low WPOI corresponding to a noninvasive growth pattern (top rows) and a high WPOI representing an invasive phenotype (bottom rows). Bright-field images of H&E-stained tongue sections were used to annotate the tumor boundaries by an expert pathologist (Fig. 5a and j). Bright-field images of adjacent, PSR-stained sections (Fig. 5b and k) were used to create collagen masks (Fig. 5c and l) by making use of the fact that collagen fibers appear red after PSR staining. PSR-stained sections were measured with ComSLI. The grayscale bright-field images of the H&E-stained sections (Fig. 5a and j) were registered onto those of the PSR-stained sections (Fig. 5b and k), in order to determine the tumor boundaries in the PSR-stained sections (peritumoral areas highlighted in blue). Figure 5e and n show the fiber orientation maps of the noninvasive and invasive tissue samples, respectively, and Fig. 5f and o the collagen fiber orientations (obtained after applying the respective collagen mask). Figure 5g and p (Fig. 5h and q) show the relative orientations of all fibers (collagen fibers) to the tumor boundary in the peritumoral regions (pixels with fibers oriented parallel or perpendicular to the tumor boundary are displayed in green or magenta, respectively).

**Fig. 5.**
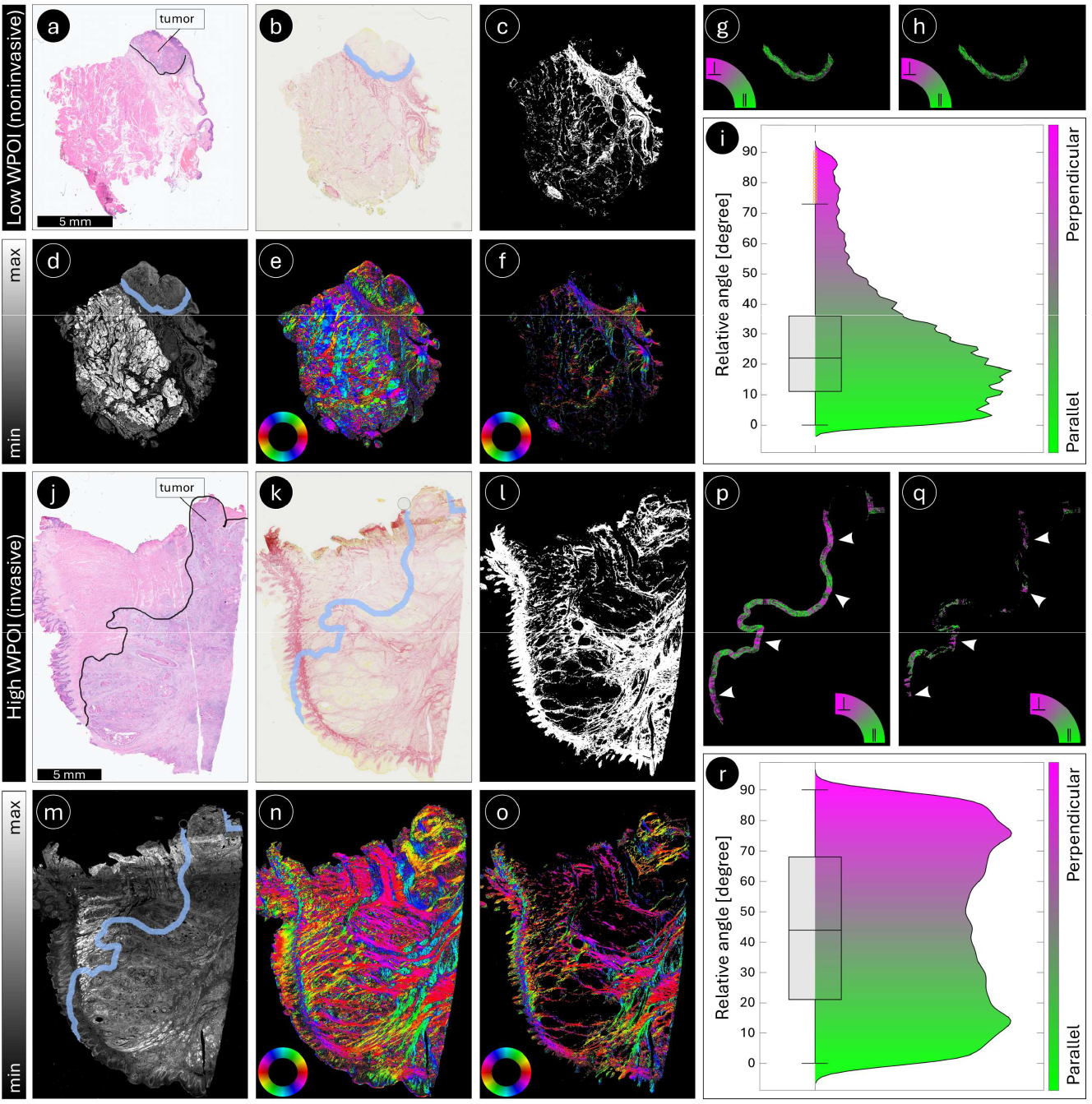
ComSLI for visualizing relative (to the tumor invasive front) fiber orientations. Top: Human tongue cancer specimen (4-μm-thin FFPE section) with a low WPOI of 2 (noninvasive tumor). Bottom: Human tongue cancer specimen (4-μm-thin FFPE section) with a high WPOI of 5 (invasive tumor). (a) and (j) Bright-field digital scans of H&E-stained sections with the tumor invasive front annotated by a solid black line. (b) and (k) Bright-field digital scans of adjacent sections stained with PSR with peritumoral areas highlighted in blue. (c) and (l) Collagen masks generated from the PSR-bright-field images by segmenting the red image pixels. (d) and (m) Average scattering maps of the PSR-stained sections with peritumoral areas highlighted in blue. (e) and (n) Corresponding fiber orientation maps, color-coded according to the color wheel in the bottom left. (f) and (o) Collagen fiber orientations obtained after applying the collagen mask. (g) and (p) Relative (to the tumor invasive front) fiber orientation maps of the peritumoral areas, color-coded according to the color arc (parallel orientations in green, perpendicular orientations in magenta). (h) and (q) Relative (to the tumor invasive front) fiber orientation maps of the peritumoral areas, masked for collagen. Arrowheads in (p) and (q) indicate passageways where tumor cells are more likely to escape, as fibers in these regions are oriented perpendicular to the tumor boundary (magenta). (i) and (r) Box and violin plots of the relative fiber orientations of collagen fibers in the peritumoral areas (as shown in (h) and (q)) for samples with low and high WPOI, respectively. As shown in the plots, the sample with a high WPOI exhibits a higher proportion of fibers oriented perpendicular (radially aligned) to the tumor invasive front compared with the sample with a low WPOI.

In the low-WPOI sample (top rows of Fig. 5), collagen fibers predominantly follow orientations that are more parallel to the tumor boundary (green). In contrast, the high-WPOI sample (bottom rows of Fig. 5) shows a marked increase in fibers aligned perpendicular to the invasive front (magenta), creating radial structures that may facilitate tumor cell dissemination (white arrowheads). Quantitative analysis of the relative orientations of peritumoral collagen fibers in Fig. 5h and q confirms these observations (see box and violin plots in Fig. 5i and r): The high-WPOI sample displays a larger fraction of collagen fibers aligned perpendicular to the tumor invasive front compared with the low-WPOI sample. These findings indicate that ComSLI captures invasion-associated remodeling of the collagen matrix and provides spatially resolved information on fiber organization linked to tumor aggressiveness.

## 4. Discussion

In this study, we explored the use of ComSLI as a whole-slide microscopy approach for characterizing fiber organization in solid tumors using routinely prepared histological sections, motivated by the need for an objective and scalable method to assess fiber architecture in archived clinical material. We applied ComSLI to a diverse set of paraffin-treated human and animal tumor samples, including glioma, colorectal cancer, and head and neck cancer, and evaluated its performance relative to polarization-based microscopy. This comparison showed that, unlike polarization-based imaging, ComSLI preserves fiber orientation contrast after standard paraffin embedding and staining. The analysis focused on the ability of ComSLI to visualize tumor growth pathways and invasion-associated stromal features, such as desmoplastic reactions, as well as collagen fiber orientations relative to the tumor invasive front. By demonstrating robust fiber orientation mapping in FFPE tissue, this work addresses a key limitation of existing fiber-imaging techniques and assesses the potential of ComSLI for large-scale retrospective studies using archived clinical material.

Together, these results position ComSLI as a powerful tool for assessing tissue microstructure across a range of biological contexts and preparation methods. Unlike birefringence-based techniques, its performance is not compromised by standard histological processing like paraffin embedding. Moreover, its sensitivity to subtle changes in fiber architecture makes it particularly well suited for studying tumor invasion and the surrounding stromal responses, including fibrosis and desmoplastic remodeling.

ComSLI is a cost-effective technique with high spatial resolution and a large field of view, enabling the detection of highly interwoven fibers and fiber crossings, independent of sample preparation. However, in its current form, ComSLI also has a few limitations.

First, the method primarily captures the orientation of in-plane fibers, while strongly inclined fibers may not be accurately resolved. Potential for estimating the out-of-plane fiber orientation from ComSLI signals has been reported in previous studies [56, 64], and a reliable reconstruction of the 3D-fiber orientation from the measured scattering signal is the subject of current research.

Second, ComSLI was used in this study in a forward-scattering configuration and applied to thin histological tissue sections. While this approach ensures direct compatibility with routine histopathology and archived FFPE material, analyzing fiber orientations in non-sectioned tissue can also be valuable, as it avoids the need for mechanical sectioning. Tissue sectioning may disrupt the native three-dimensional continuity of fibrous networks. Back-scattered light imaging approaches applied to intact tissue surfaces prior to sectioning can provide information on fiber organization without mechanical disruption and may enable assessment of larger-scale structural heterogeneity. Extending ComSLI to back-scattered light measurements could therefore offer a more comprehensive characterization of tumor-associated fiber architecture and represents an interesting direction for future work.

Third, ComSLI is not specific to collagen. It visualizes all kinds of fibrous structures, including muscle fibers (cf. Fig. 5). As a result, when studying the organization of collagen fibers in cancer samples, collagen-specific staining, such as PSR or trichrome, is currently required to exclude non-collagenous fibers. However, in clinical practice, samples are typically stained with H&E to identify the tumor margins (which is why we needed to register the H&E tumor margins onto the PSR-adjacent sections, see Fig. 5). In the future, ComSLI could be combined with virtual collagen staining approaches to achieve collagen specificity also in H&E-stained sections, including deep learning-based analysis of bright-field images [71], spectral phasor analysis of eosin fluorescence images [72] or hyperspectral imaging of H&E-stained sections [73].

So far, the role of fiber directionality in certain cancers, such as OSCC, has not been studied extensively. Previous analyses have relied on time-consuming and expensive techniques that cannot be applied to whole-slide FFPE archived sections, substantially limiting the size of study cohorts. To investigate the role and clinical relevance of collagen fiber re-alignment in the progression of oral cancer, two elements are required: (I) an objective, fast, inexpensive, and non-laborious technique to quantify this process, and (II) a large number of tumor samples linked to clinical follow-up data.

The current study is a proof-of-principle study, which is focused on demonstrating the performance and utility of ComSLI across different cancer types and histological preparations. As follow up, we aim to: (I) use ComSLI to interrogate the complex interwoven microscopic organization of collagen fibers in archived FFPE sections at a macroscopic scale with micrometer resolution, and (II) utilize a recently established OSCC database, RONCDOC [74], one of the largest and most detailed OSCC patient databases worldwide.

ComSLI is a label-independent technique that enables high-resolution, large-scale imaging of archived tissue architecture. It provides a fast, cost-effective, and objective method for evaluating histopathological features. The fiber orientation extraction workflows are automated and based on pre-validated algorithms, which suggests that clinical implementation could be straightforward.

## 5. Conclusion

This study demonstrates that computational scattered light imaging (ComSLI) can be applied to visualize fiber organization in solid tumors with high resolution and across large fields of view. The ability of ComSLI to directly quantify fiber orientations, even in complex tissues with multiple fiber crossings, provides a distinct advantage over existing imaging modalities while maintaining compatibility with standard FFPE tissue preparations. Previous studies applied ComSLI to (diseased) brain tissue and healthy non-brain tissue. By extending ComSLI beyond its established applications in neural tissue to tumor histopathology, we highlight its potential as an objective, label-free tool for evaluating collagen remodeling, a process closely linked to tumor aggressiveness and metastatic potential. Given the consistent association between collagen fiber reorganization and patient outcomes across multiple cancer types, ComSLI could serve as a powerful prognosticator of occult metastasis, disease recurrence, and survival. Ultimately, integrating this technique into clinical workflows may provide pathologists and oncologists with actionable information for risk stratification and treatment planning, thereby advancing more personalized cancer care.

## Author contributions

H.A. contributed to the study design, performed the ComSLI measurements, analyzed and interpreted the ComSLI data, wrote the first draft of the manuscript, and prepared the figures. L.E. contributed to the data interpretation. R.V.E. performed the PLI measurements and analysis. M.E. developed code to calculate fiber orientations relative to the tumor boundary, created the collagen masks, and registered the bright-field images. M.D. annotated the rectal tissue sample and interpreted the observed fiber orientations using histological images. S.A.K. annotated the oral tissue sample and interpreted the observed fiber orientations using histological images. S.K. contributed to the study design, data interpretation, and supervision. M.M. contributed to the study design, data interpretation, supervision, and manuscript editing. All authors reviewed the manuscript.

## Acknowledgement

The authors would like to thank Markus Cremer and the laboratory team at Forschungzentrum Jülich (INM-1), Germany, for providing the rat brain section, and Meedie Ali and Laura Mezzanotte (Department of Radiology and Nuclear Medicine, Erasmus MC Cancer Institute) for providing mouse glioma sections. The authors would like to further express their gratitude to Jose A. Hardillo (Department of Otorhinolaryngology, Head and Neck Surgery, Erasmus MC Cancer Institute) for providing clinical insight to the study.

## Supplemental document

See Supplement 1 for supporting content.

## Supplemental Document

**Fig. S1.**
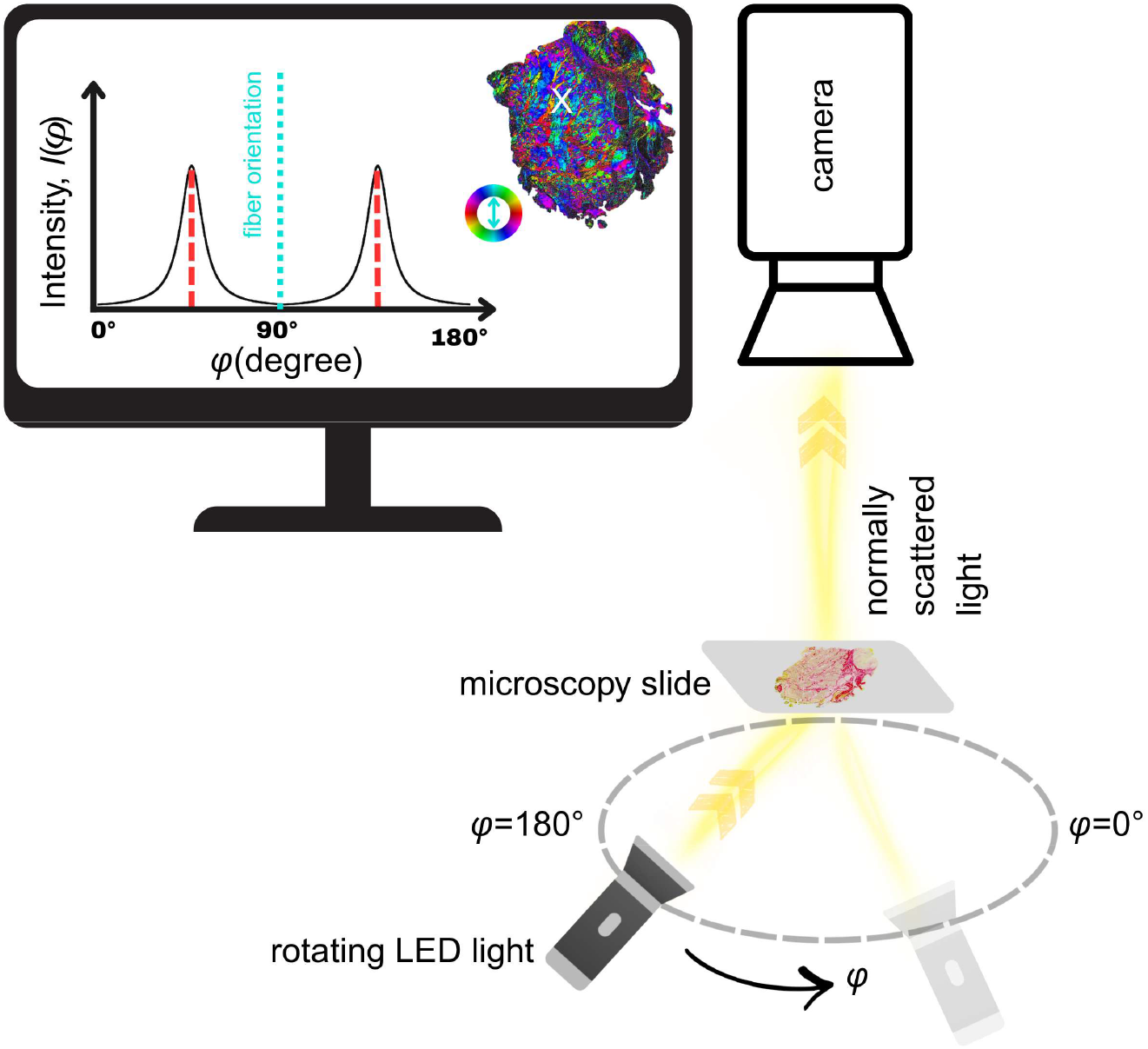
ComSLI setup. A rotating LED lamp illuminates the tissue section from below under a fixed polar angle of about 45°. A camera records the normally scattered light for different rotation angles. The intensity of each image pixel in the resulting image series, *I*(*φ*), is evaluated. The monitor shows an exemplary intensity profile of one image pixel; the in-plane fiber orientation is determined by the mid-position of a peak pair and displayed according to a color wheel (here, the fiber orientation is *φ*=90° and shown in cyan).

**Fig. S2.**
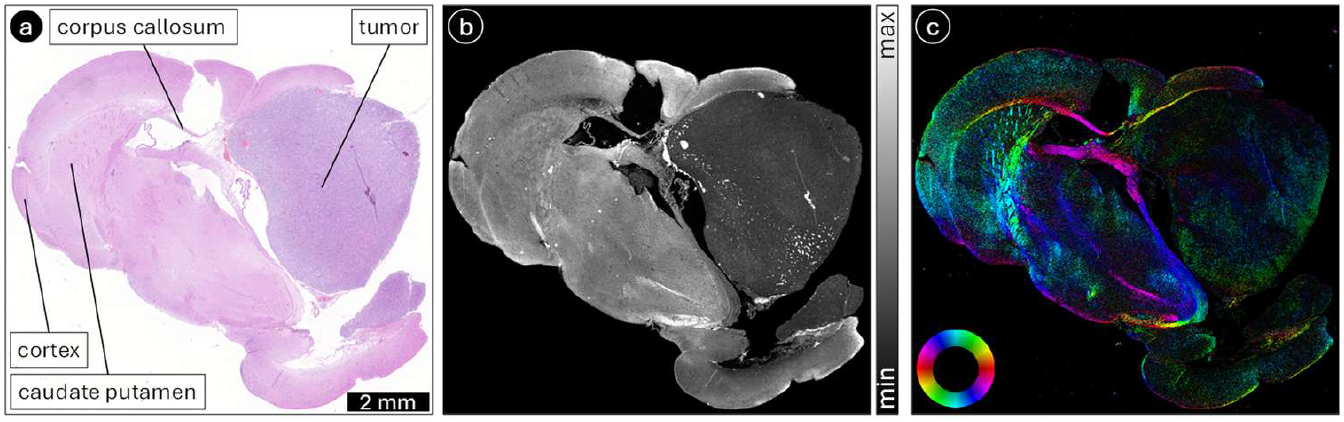
ComSLI in an additional mouse glioma brain section (H&E-stained, 4-μm-thin, FFPE). (a) Bright-field image with tumor and selected anatomical regions labeled. (b) Average scattering map. (c) Fiber orientation map.

